# The role of the gut microbiota in the effects of early-life stress and dietary fatty acids on later-life central and metabolic outcomes in mice

**DOI:** 10.1101/2021.11.02.467036

**Authors:** Kitty Reemst, Sebastian Tims, Kit-Yi Yam, Mona Mischke, Jan Knol, Stanley Brul, Lidewij Schipper, Aniko Korosi

## Abstract

Early-life stress (ELS) leads to increased vulnerability for mental and metabolic disorders. We have previously shown that dietary low ω-6/ω-3 polyunsaturated fatty acid (PUFA) ratio is able to protect against ELS-induced cognitive impairments. Due to the importance of the gut microbiota as determinants of long-term health, we here study the impact of ELS and dietary PUFA’s on the gut microbiota, and how this relates to the previously described cognitive, metabolic and fatty acid profiles.

Male mice were exposed to ELS via the limited bedding and nesting paradigm (postnatal day (P)2 – P9) and to an early diet (P2 – P42) with either high (15) or low (1) ω-6 linoleic acid to ω-3 alpha-linolenic acid ratio. 16S ribosomal RNA was sequenced and analyzed from fecal samples at P21, P42 and P180.

ELS increased β-diversity at P42, which persisted into adulthood. The low ω-6/ω-3 diet prevented the ELS-induced increase in β-diversity, at P42. At the level of taxa abundance, for example, the abundance of the phyla Bacteroidetes increased while Actinobacteria and Verrucomicrobia decreased with age; ELS reduced the relative abundance of the genera *RC9 gut group* and *Rikenella* into adulthood and the low ω-6/ω-3 diet reduced the abundance of the Firmicutes Erysipelotrichia. At P42, species abundance correlated with body fat mass and circulating leptin (e.g. Bacteroidetes and Proteobacteria taxa) and fatty acid profiles (e.g. Firmicutes taxa).

This study gives novel insights into the impact of age, ELS and dietary PUFAs on microbiota composition, providing potential targets for non-invasive (nutritional) modulation of the ELS-induced deficits.

**Importance:** Early-life stress (ELS) leads to increased vulnerability to develop mental and metabolic disorders, however the biological mechanisms leading to such programming are not fully clear. Increased attention has been given to the importance of the gut microbiota as determinant of long term health and as potential target for non-invasive nutritional strategies to protect against the negative impact of ELS. Here we give novel insights in the complex interaction between ELS, early dietary ω-3 availability and the gut microbiota across ages and provides new potential targets for (nutritional) modulation of the long-term effects of the early-life environment via the microbiota.

## Introduction

There is ample clinical and preclinical evidence that early-life stress (ELS) is associated with increased vulnerability to mental and metabolic health problems such as depression and inflammatory bowel disease^1–4^. We and others have shown in recent years that chronic ELS induced in rodent models via the limited bedding and nesting material (LBN) paradigm^5,6^ leads to impaired cognitive functions and an altered metabolic profile^7,8^. Moreover, we demonstrated that early postnatal exposure to diet with a low ω-6 to ω-3 polyunsaturated fatty acid (PUFA) ratio was able to protect against the ELS-induced cognitive deficits without affecting the metabolic alterations^9^. Currently the exact underlying mechanisms for the effects of ELS and the beneficial effect of the diet are not fully understood and may be multi-factorial. In this paper we address the effects of ELS and dietary ω-6/ω-3 PUFA ratio on fecal microbiota and if and how these relate to the effects of ELS and early postnatal diet on both the brain and metabolism across different ages that we reported earlier^9^.

In recent years, there has been an increasing interest in how the gut microbiome might impact our health^10,11^. Particular attention has been devoted to the cross talk between the gut microbiota and the brain, known as the microbiota – gut – brain (MGB) axis, an integrated communication system including neural, hormonal and immunological signaling pathways through which the gut microbiota can influence brain development and function and vice versa^12,13^. Increasing evidence supports the intriguing hypothesis that the microbiota can influence brain functions, that dysbiosis might contribute to changes in behaviour (e.g. social behaviour^14^) and the development and etiology of brain disorders (e.g. depression^15–18^) and that targeting the microbiota is effective in modulating brain function (e.g. cognitive functions^19^). Similarly, the gut microbiota are also thought to impact greatly on the immune system and metabolic health and has been associated with various risk factors of obesity and metabolic syndrome^20^.

Several elements are emerging to be key in modulating the microbiome composition, including developmental life stages, stress and diet^13,21,22^. In fact, the development of the microbiome coincides with crucial (neuro)developmental periods. While little is known of the exact developmental trajectory of the microbiome in mice, we know from human literature that the intestinal microbiome starts to develop during and shortly after birth, during which time the brain is also going through immense developmental changes^23^. Preclinical evidence shows that various early postnatal stress paradigms, in different species, impact the gut microbiota^24^. For example, maternal separation (MS) has been shown to increase intestinal permeability in rats^4,25^, and affect the microbiota composition of the gut microbiota of infant Rhesus monkeys directly after separation^26^ and of 7-week old rats^27^. Such microbial composition changes may be instrumental for establishment of some of the MS-induced anxiety-related alterations, as germ free mice were not affected by MS to the same extend as colonized mice^28^. Also chronic ELS induced via the limited bedding and nesting material (LBN) paradigm^6^ in male rats led to changes in microbiota composition and increased intestinal permeability at weaning age^29^. Thus the early-life adversity-induced dysbiosis could possibly contribute to later life mental and metabolic health^10,23,24,30,31^.

Next to development and exposure to early adversity, diet, and more specifically, dietary PUFA composition has also been shown to modulate the composition of the gut microbiota at different stages of life^32,33^. For example, an 8-week supplementation with ω-3 long chain (LC)PUFAs including docosahexaenoic acid (DHA) and eicosapentaenoic acid (EPA) in middle-aged healthy individuals led to multiple changes in bacterial taxa including an increased abundance of genera involved in butyrate production, which have been suggested to be important for mental health^15,34^. The abundance of dietary ω-3 and ω-6 PUFAs during early life phases is also highly relevant as these are key factors for proper development and function of the brain^35^ and can influence the microbiome^33^.

In the last century, there has been a marked change in the consumption of ω-6 and ω-3 PUFAs, with a high intake of especially ω-6 linoleic acid (LA) in western societies, resulting in a high ω-6/ω-3 ratio^36^. Given the relevance of dietary ω-6/ω-3 for brain development and function^35^, this shift is thought to increase today’s prevalence of psychopathology and chronic disease^37^, and possibly also contributes to gut dysbiosis and thereby impacting on the MGB-axis^32^. Therefore, dietary fatty acids have been explored as possible strategy to modulate (stress-induced) behavioral changes and cognitive functioning^9,38–41^. In particular, the possible protective actions of ω-3 PUFA during different life stages on the early-life stress induced effects have been explored^9,40,42^. Pusceddu and colleagues demonstrated that long-term exposure to a diet with low ω-6/ω-3 ratio (i.e. between 5 and 17 weeks of age by supplementation with ω-3 LCPUFAs including DHA) was beneficial for anxiety and cognition in non-stressed female rats, and could restore part of the disturbed gut-microbiota composition of MS female rats which was associated with the attenuation of the cortisol response to an acute stressor^40,42^. While this consisted of a lifelong intervention starting at 5 weeks of age, we have recently shown that a relatively short dietary intervention with low ω-6/ω-3 diet starting in the early postnatal period (i.e. from postnatal day 2 - 42), is able to restore the effects of ELS (via LBN) exposure, on brain FA composition early in life and on cognitive functions and brain plasticity in adulthood, without modulating the ELS-induced alterations in body fat mass and circulating leptin in mice^9^.

Here we study the effects of ELS, using the LBN paradigm in mice (P2 and P9), an early dietary intervention with low ω-6 linoleic acid (LA) to ω-3 alpha-linolenic acid (ALA) ratio (P2 and P42) and their interaction on the short-term (at P42) and long-term (at P180, after exposure to regular diet from P42 onwards) impact on the gut microbiota composition and if and how these changes relate to the earlier reported central and metabolic ELS-induced profiles described in the same cohort of mice^9^.

## Material and Methods

### 2.1 Animals

In the current study we describe microbiome data from the same mice from our previous publication ^9^. In brief, male (6 weeks old) and primiparous female (8 weeks old) C57Bl/6J mice were purchased from Harlan Laboratories B.V. (Venray, the Netherlands). After arrival at the animal facility, the mice were put on a synthetic AIN-93G diet (Ssniff-Spezialdiäten GmbH, Germany)^43^ and housed in a controlled environment (temperature 22±1°C, humidity 55±5%) with *ad libitum* food and water, under a 12:12 h light-dark cycle schedule (lights on at 8 AM). After two weeks of acclimatization, mice were bred in house by housing two females with one male for one week in a type-II long cage. Subsequently females were housed in single-sex pairs for another week, and after that pregnant females were housed individually in a standard cage (type-I short cage) covered with a filter top. Females were monitored daily, between 9 and 10 AM, for the birth of pups. When a litter was detected, the previous day was designated the day of birth (postnatal day (P)0). At P2, dams with litter were randomly assigned to control (CTR) or ELS condition, see 2.2 and to one of the experimental diets (see 2.3). At P21, offspring was weaned and male offspring was housed in groups (littermates; 2 or 3 animals per cage) in type-II long cages with standard amount of bedding material. Mice were kept on their respective diet until P42, after which all groups were switched to standard semi-synthetic diet (AIN93M)^43^ until end of the study.

All experimental procedures were approved by the Animal Welfare Body of the University of Amsterdam, and the Central Authority for Scientific Procedures on Animals (CCD – Centrale Commissie Dierproeven) in compliance with Dutch legislation and the principles of good laboratory animal care following the EU directive for the protection of animals used for scientific purposes.

### 2.2 Chronic early-life stress exposure

We used the chronic ELS model, based on the limited bedding and nesting (LBN) stress paradigm as described before by our group and others^5,7,9^. The LBN paradigm induces fragmentation of maternal care which results in chronic stress in the pups. At P2, litters were culled to six pups per litter (sex ratio m:f of 3:3 or 4:2) without cross fostering, randomly assigned to CTR or ELS condition. In ELS cages, the bottom was covered with a little amount of sawdust bedding and a fine-gauge stainless steel mesh is placed 1 cm above the cage floor. Half a square piece of cotton nesting material (2,5 x 5 cm, Technilab-BMI, Someren, the Netherlands) was placed on top of the mesh. Control cages were equipped with standard amounts of sawdust bedding and nesting material (one square piece of cotton nesting material (5×5cm). Cages were equipped with food and water *ad libitum* and covered with a filtertop. Throughout all procedures, manipulation was kept to a minimum to avoid handling effects and animals were left undisturbed until P9. On the morning of P9, bodyweight of the dams and pups and the consumed amount of food and/or water was measured, this data can be found in our previous publication^9^. From P9 onwards all animals were housed in cages equipped with a standard amount of nesting and bedding material.

### 2.3 Experimental diets

Experimental diets were provided from P2 onwards to dam with litter, and after weaning (P21) offspring were kept on their respective diet until P42. During lactation, fatty acid composition of the maternal diet, in particular LA and DHA, is reflected in milk fatty acid composition^43^. The two experimental diets (Ssniff-Spezialdiäten GmbH, Soest, Germany) were semi-synthetic differing only in LA/ALA ratio that was either a high (15) or low (1.1) The diets were isocaloric and contained a macro- and micronutrient composition according to the AIN93-G purified diets for laboratory rodents^44^ (Supplementary Table S1). Experimental groups are referred to as: CTL- and ELS-high; control and ELS-exposed animals that received a diet with ω-6/ω-3 ratio=15 and CTL- and ELS-low; control and ELS-exposed animals that received a diet with ω-6/ω-3 ratio=1.1.

### 2.4 Fecal sample collection, DNA extraction and sequencing

Fresh fecal samples were collected during a brief handling moment of approximately 2 minutes, from three separate age cohorts, P21, P42 and P180. One or two pellets per animal were snap frozen and stored at −80°C until further analysis. The N per group was as follows: P21: CTL-high: 3, ELS-high: 5, CTL-low: 5, ELS-low: 5; P42: CTL-high:9, ELS-high: 14, CTL-low: 7, ELS-low: 7; P180: CTL-high: 11, ELS-high: 11, CTL-low: 10, ELS-low: 9.

DNA extraction from these samples was performed with QIAmp DNA Stool Mini Kit (Qiagen) according to the manufacturer’s protocol except for the addition of two bead-beating steps. To 0.2 – 0.3 g of fecal sample 300 mg of 0.1 mm glass beads together with 1.4 mL of ASL (lysis) buffer and on this suspension the first bead-beating step was applied for 3x 30 sec (FastPrep-24 instrument program 5.5). After addition of the InhibitEx tablet the second bead-beating step was applied for 3x 30 sec (FastPrep-24 instrument program 5.5) to homogenize the sample. Following each bead-beating step samples were cooled for 5 min on ice. Extracted DNA purity was checked using the NanoDrop™ spectrophotometer (Thermo Fisher Scientific Inc.), whereas DNA quality and concentration was measured using the Quant-iTTM 193 dsDNA BR Assay kit (Invitrogen™). DNA aliquots were stored at −20°C until use.

On the purified fecal DNA extracts primers Bact-0341F (5′-CCTACGGGNGGCWGCAG-3′) and Bact-0785R (5′-GACTACHVGGGTATCTAATCC-3’)^45^ were used to amplify the V3–V4 regions of the bacterial 16S rRNA gene and the generated amplicons were subsequently sequenced on a Illumina MiSeq instrument as described previously^46^.

### 2.5 Sequencing analysis

Sequencing data was analyzed using the Quantitative Insights Into Microbial Ecology (QIIME) v.1.9.0 pipeline^47^. Sequences with mismatched primers were discarded. Quality control filters were set to retain sequences with: a length between 200 and 1000 bases; a mean sequence quality score >15 in a five-nucleotide window; no ambiguous bases. The filtered sequences were grouped into Operational Taxonomic Units (OTUs) by *de novo* OTU picking using the USEARCH algorithm^48^ at 97% sequence identity. Subsequently, the Ribosomal Database Project Classifier (RDP)^49^ was applied to assign taxonomy to the representative sequence (i.e. the most abundant sequence) of each OTU by alignment to the SILVA ribosomal RNA database (release version 1.1.9)^50^. ChimeraSlayer^51^ was applied, as part of QIIME, to filter for chimeric sequences and these were excluded from all downstream analyses. Representative OTUs were aligned using PyNAST^47^ and used to build a phylogenetic tree with FastTree^52^, which was used to calculate UniFrac^53^. OTUs that could not be aligned with PyNAST, singletons and low abundant OTUs with a relative abundance <0.002% were excluded. Weighted UniFrac distances were used to assess the (dis)similarities between the samples^53,54^. Rarefaction was applied to the OTUs by QIIME to ensure identical number of reads per sample in order to perform α-diversity calculations using the Chao1 metric.

### 2.6 Statistical analyses

The microbial diversity within each sample (α-diversity) was assessed to investigate the overall microbiota development between P21, P42 and P180. The average Chao1 value for each sample at each sequencing depth, resulting from the rarefaction procedure, was visualized. The average Chao1 values were grouped per diet, condition, as per age and expressed as mean ± standard error of the mean (SEM). The Three-Way Generalized Linear Mixed Model (GLMM) was performed at the a sequencing depth of 11,535 sequences. Considering the low sample sizes of the P21 samples (n = 3 to 5 per experimental group), these were excluded from further analyses. Therefore, a Two-Way GLMM was performed only at P42 and P180 separately, at the highest sequencing depth.

Between sample microbiota profile (dis)similarity (β-diversity), was assessed both on OTU level with Weighted Unifrac and on genus level, with aggregated data, at P42 and P180 separately (due to the strong separation between the two ages (Supplementary Fig. 1A). At each age a Two-Way ANOVA with two predictor variables (i.e., condition and diet) and interactions thereof on the Weighted Unifrac distances within the four experimental groups (homogeneity) was performed. In addition, Weighted Unifrac distances of samples between the four experimental groups were plotted and analyzed with a Kruskal-Wallis analysis of variance and Dunn’s multiple comparison test (Supplementary Fig. 1B. Principal component analysis (PCA) and distance-based redundancy analysis (db-RDA), using Bray-Curtis metrics, were performed on the OTU data aggregated at genus-level taxonomy to assess the influence of condition, diet and their interactions on the fecal microbiota composition at each age separately. Since litter effects have been shown to drive gut microbiota variation in common laboratory mice^55^, litter correction was applied to the db-RDA calculations. Data at genus level was Log transformed and standardized by Hellinger transformation^56^. Significance of the explained variance in the db-RDAs were assessed with ANOVA-like permutation test for Redundancy Analysis^57^. The 10 genera explaining the most variation in the PCA and db-RDA were visualized. The db-RDA procedures were performed using the vegan package(version2.5-7) in R(version 3.6.2).

Next, the impact of condition, diet, and age on the microbial taxa abundances was investigated. To this end the sequence data was aggregated at the following taxonomic levels: genus, family, order, and at phylum. Also for microbial taxa abundances, litter was accounted for in the statistical analysis. GLMM was used to determine whether litter has a significant effect on the sequence data derived abundances of a taxonomic group within every taxonomic level and in cases where it did, it was taken along as a co-variate in the GLMM. In order to estimate the effect of age, a Three-Way GLMM was performed on the sequence data derived abundances of each taxonomic group within every taxonomic level, and in order to estimate the effect of condition and diet, a Two-Way GLMM was employed on each age group (P42 and P180) separately. After performing an GLMM, the resulting sets of p-values, one set for each of the predictor variables and interactions thereof, were used to estimate the false discovery rate (FDR) by calculating q-values^58^. Resulting p-values <0.05 with corresponding q-values <0.1 were regarded as significant.

#### Correlational analysis between abundance of microbial species and central and peripheral outcomes

Finally, using a Spearman correlation test we tested whether the abundance of microbial species on four taxonomic levels from the current study correlated with previously reported parameters from the same mice^9^. The parameters were: cognitive behaviour (Performance on Object Location Task and Morris Water Maze), fatty acid profiles in hippocampus, liver and erythrocytes and multiple metabolic outcomes (bodyweight, plasma leptin levels, inguinal fat, sum white fat). Detailed description of the methods regarding these parameters can be found in Yam et al. (2019). All correlations are shown in Figure 5, and correlations with ρ>0.7 are reported in the text and Supplementary Table S2 and S3.

## Results

### 3.1 Low dietary ω-6/ω-3 ratio prevents the early-life stress induced expansion of microbiota diversity in fecal samples

Male mice were exposed to ELS via LBN paradigm (P2 to P9) and to an early diet (P2 – P42) with either high (15) or low (1) ω-6 linoleic acid to ω-3 alpha-linolenic acid ratio, and are the same cohort of mice as in our previous publication^9^. 16S ribosomal RNA was sequenced and analyzed from fecal samples at P21, P42 and P180 (Fig. 1A). Experimental groups are referred to as: CTL- and ELS-high; control and ELS-exposed animals that received a diet with ω-6/ω-3 ratio=15 and CTL- and ELS-low; control and ELS-exposed animals that received a diet with ω-6/ω-3 ratio=1.1 (Methods).

**Figure 1.**
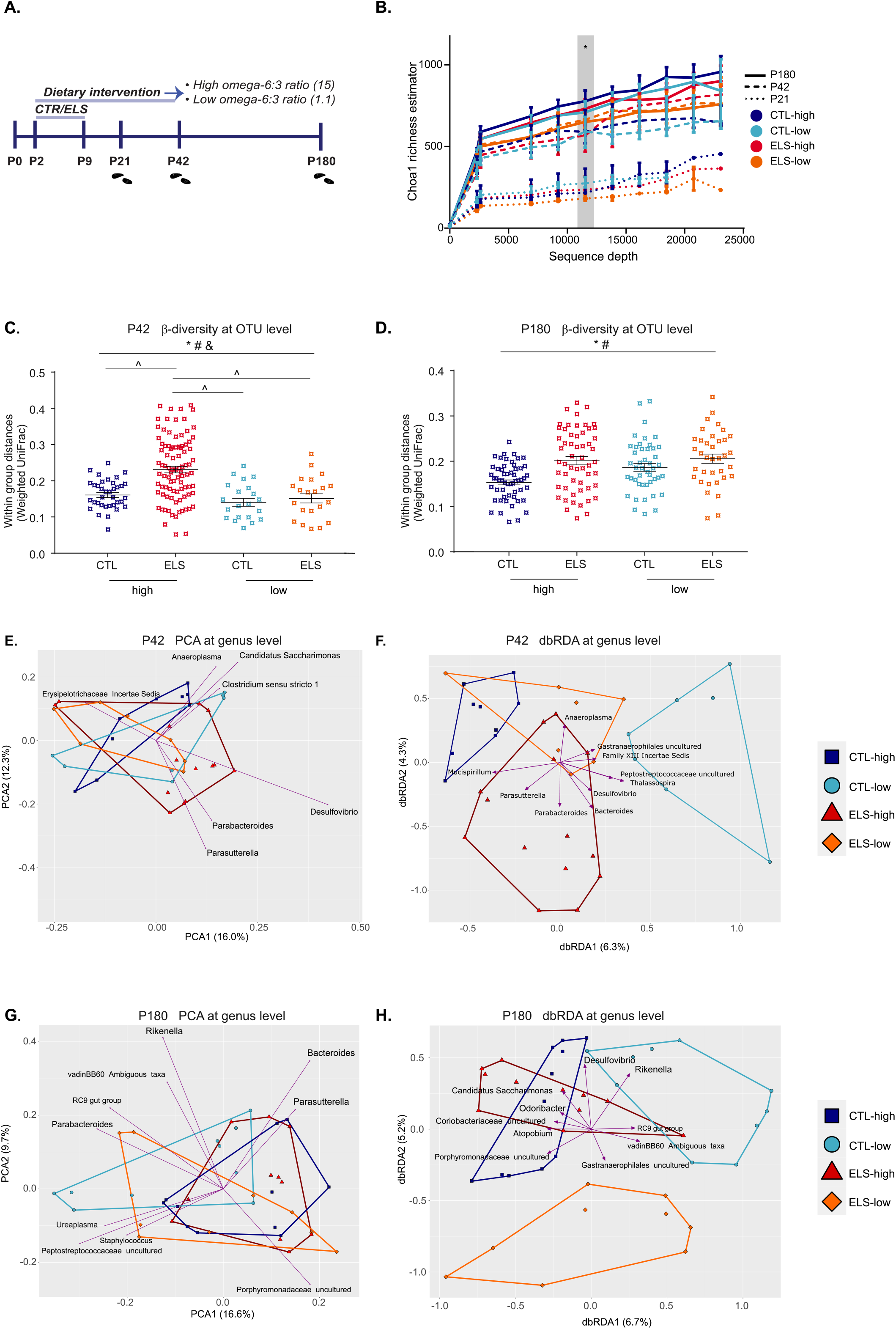
Early dietary low ω-6/ω-3 diet reverses ELS-induced increase in microbiota β-diversity. **A:** Experimental timeline. **B:** Chao1 plot displaying increase in α-diversity with age (GLMM at all sequencing depths p<0.0001; all experimental samples present at a sequencing depth of 11,535 sequences). **C, D:** Average weighted UniFrac distances (within groups) of β-diversity on OTU level comparing phylogenetic configurations of fecal microbial communities of the four experimental groups for both ages, Two-Way ANOVA; *: condition effect, #: diet effect, &: interaction effect condition*diet, ^: significant difference with Tukey *post-hoc* test. P<0.05. **C:** P42, showing the increase in β-diversity in ELS-high and not in ELS-low experimental group. **D:** P180, showing increase in β-diversity in ELS-high and in ELS-low experimental group. **E, F, G, H:** Principal Component Analysis (PCA) and distance based Redundancy Analysis (db-RDA) of β-diversity aggregated at genus level for both ages. The 10 genera explaining most variation in the PCA and db-RDA were visualized. **E:** PCA at P42. **F:** db-RDA at P42, ANOVA like permutation test for Redundancy Analysis, p=0.018. **G:** PCA at P180. **H:** db-RDA at P180, ANOVA like permutation test for Redundancy Analysis, p=0.003). Abbreviations: GLMM: General Linear Mixed Model. OTU: Operational Taxonomic Unit. PCA: Principal Component Analysis. Db-RDA: Distance-based Redundancy Analysis

#### α-diversity

*α-diversity* summarizes the distribution of taxa abundances in a given sample into a single number that depends both on species richness and evenness. For all four experimental groups, the lowest Operational Taxonomic Unit (OTU) level *α*-diversity within samples was observed at P21 and increased with age (GLMM: *timepoint* p<0.0001 at a sequencing depth of 11,535 sequences; all experimental samples present at this sequencing depth) (Fig. 1B). No differences were seen in phylogenetic *α*-diversity between the four experimental groups at any timepoint (P21, *condition*diet* F_1,14_ =0.690, P=0.420; P42, *condition*diet* F_1,33_ =0.389, P=0.537; P180, *condition*diet* F_1,37_ =0.668, P=0.419).

Our sample size at P21 was relatively low (N=3-5 per group), and even though our methodology was able to pick up age-related changes in α-diversity for further outcome measurements we only analyzed the P42 and P180 time points.

#### β-diversity

Where alpha diversity focuses on community variation within a community (sample), *β*-diversity quantifies (dis-)similarities in microbiota composition between samples. Analysis of the *β*-diversity at OTU level between samples within the experimental groups showed a main effect of condition and diet at both P42 and P180, and an interaction effect between condition and diet at P42 (Two-way ANOVA: P42, *condition*: F_1,165_ = 9.425, P = 0.0025; *diet*: F_1,165_=14.17, P=0.0002, *condition*diet* F_1,165_=5.063, P=0.0258; P180: *condition*: F_1,187_ =16.08, P < 0.0001; *diet* F_1,187_=5.002, P=0.0265). Further post hoc testing revealed that at P42, the microbiome of animals exposed to ELS and high dietary ω-6/ω-3 (ELS-high) animals displayed an increase in β-diversity as compared to the other P42-age groups (Tukey post-hoc test: ELS-high – CTL-high, P < 0.0001; ELS-high – CTL-low, P < 0.0001; ELS-high – ELS-low, P < 0.0001) (Fig. 1C,D).

Kruskal Wallis analysis of variance of the β-diversity *between* experimental groups at P42 and P180 was significant (Kruskal Wallis p<0.0001 for both ages) (Supplementary Fig. 1C,D). Dunn’s multiple comparison test at P42 showed that microbiota composition profile of “CTL-high versus ELS-low” is less distant than “CTL-high versus ELS-high” (Dunn’s post hoc: p=0.0001). The composition profile of “CTL-high versus ELS-high” is more distant than “CTL-low versus CTL-high” (p=0.0080) and “CTL-low versus ELS-low” (Dunn’s post hoc: p=0.0022. The microbiota composition of “ELS-low versus ELS-high” is also more distant than that of “CTL-low versus CTL-high” (p=0.0178) and “CTL-low versus ELS-low” (0.0050). Dunn’s multiple comparison test at P180 showed that the composition of “CTL-high versus ELS-low” was significantly less distant than “ELS-low versus ELS-high” (p=0.0249) and “CTL-low versus ELS-low” (p=0.0011). Next, the microbiota composition of “CTL-high versus ELS-high” was less distant than “ELS-low versus ELS-high” (p=0.0040), “CTL-low versus ELS-low” (p=0.0001) and “CTL-low versus ELS-high” (p=0.0457). Lastly, the microbiota composition of “CTL-low versus CTL-high” was less distant than “CTL-low versus ELS-low”.

Assessment of the β-diversity at genus level by principal component analysis (PCA) showed no distinct clustering of the four experimental groups at P42 (Fig. 1E), nor at P180 (Fig. 1G). Similar to the Weighted UniFrac analysis, the ELS-high group showed the lowest homogeneity (i.e., the samples of ELS-high displayed the largest spread over the plot) at P42. When performing distance-based redundancy analysis (db-RDA), distinct clustering of the experimental groups was observed at P42 and P180 (Fig. 1F,H). At P42 the condition*diet interaction explained 12,8% of the total variation (with 10,6% in the first two db-RDA axes, see Fig. 1F), this was found to be significant (ANOVA like permutation test for Redundancy Analysis; p=0.018). For P180 the condition*diet interaction explained 13,9% of the total variation (with 11,9% in the first two db-RDA axes, see Fig. 1H) which was found to be significant (ANOVA like permutation test for Redundancy Analysis p=0.003).

### 3.2 Fecal microbiota composition is affected by age, early-life stress and the ω-6/ω-3 PUFA ratio of an early diet

Analysis of relative abundances at phylum, class, family and genus levels shows that the fecal microbiota composition of the experimental groups differed significantly for several bacterial taxa at both P42 (Fig. 2) and P180 (Fig. 3). All statistical differences are included in Table 2 and Table 3, and additional descriptive information on all measured bacterial species stratified per taxonomic level, age and experimental group are included in supplementary Table S4. Analysis at phylum level indicated that for both ages the fecal microbiota was dominated by three major phyla: Bacteroidetes, Firmicutes, and Verrucomicrobia, but also Proteobacteria, Deferribacteres and Actinobacteria were present (Fig. 2A). Other phyla detected in low abundance (<3%) were Candidate division TM7 (2.1%), Cyanobacteria (0.67%) and Tenericutes (1.53%) (not depicted in Fig. 2A). Analysis at genus level showed that the twenty most abundant genera for both ages were *Parasutterella*, *Bacteroides, Atopobium*, *Bilophila, Desulfovibrio, Allobaculum, Lachnospiraceae_uncultured* and *Blautia*, *Odoribacter, Alloprevotella, Alistipes, RC9-gut group, Rikenella, Ruminococcaceae_uncultured, Anaerotruncus* and *Incertae-sedis, S24-7_uncultured, VadinBB60_uncultured, Akkermansia* and a group with unassigned sequences (Fig. 2B).

**Figure 2.**
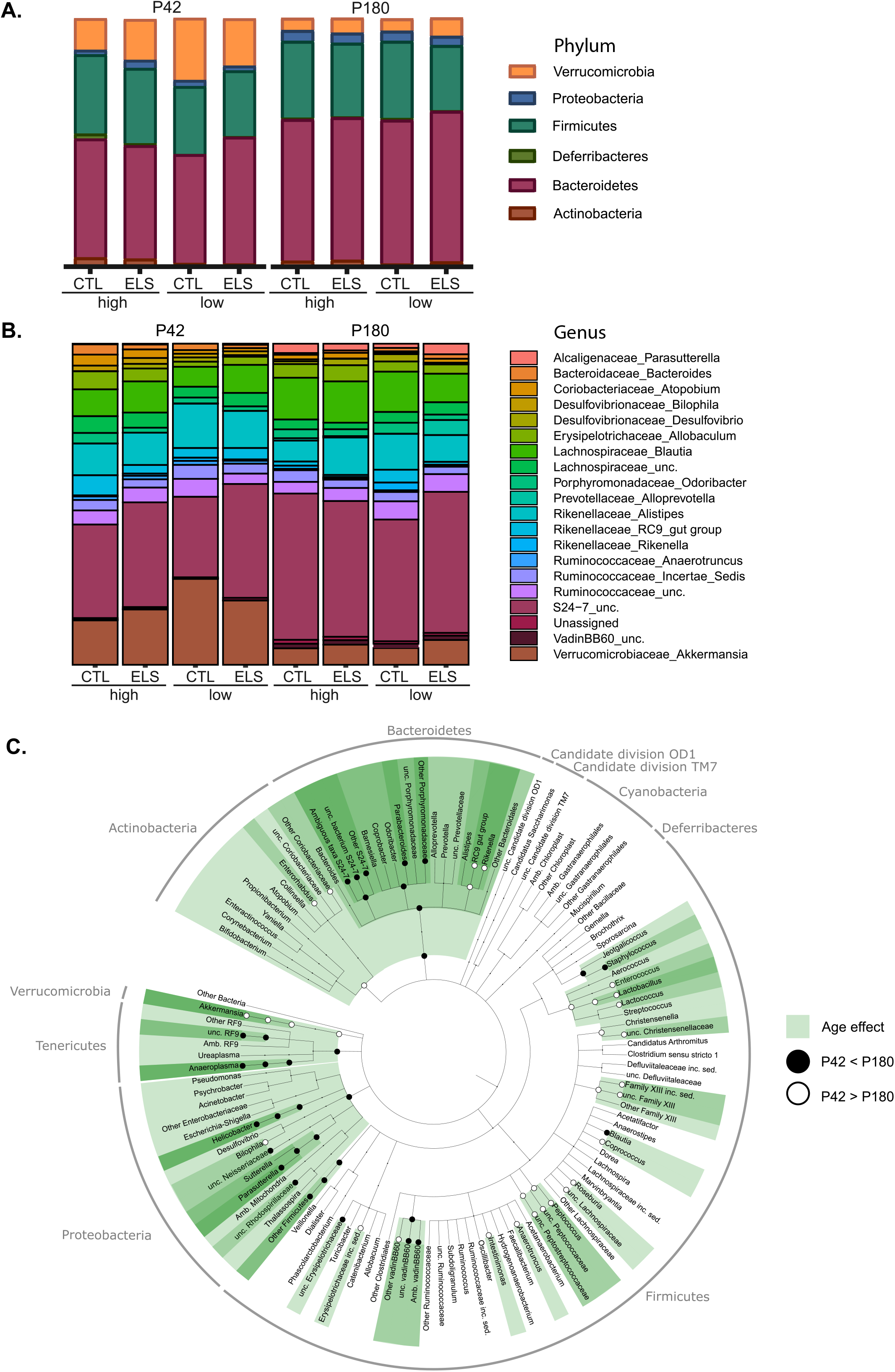
Microbiota composition goes through large amount of changes between P42 and P180. **A:** Relative abundance at phylum level for P42 and P180. **B:** Relative abundance at genus level for P42 and P180. **C:** Cladogram showing significant age-mediated changes in relative abundance of bacterial species at several taxonomic levels.

**Figure 3.**
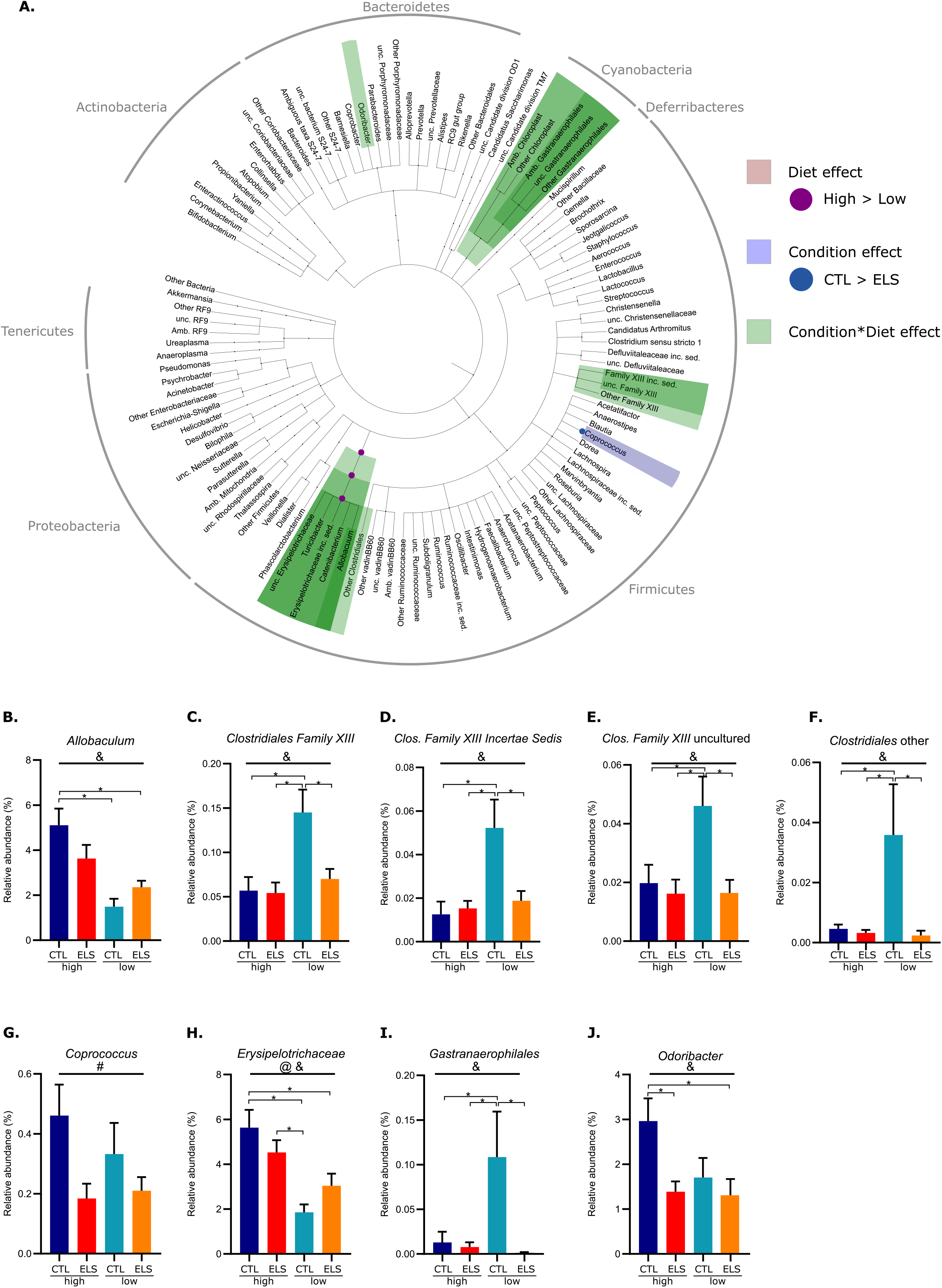
Early-life stress and early dietary ω-6/ω-3 ratio affect the microbiota composition at P42 in interaction with each other. **A:** Cladogram showing significant condition and diet-mediated changes in the relative abundance of bacterial taxa at several taxonomic levels at P42. **B – J:** Bar graphs of detected interaction effects (condition*diet) for bacterial taxa at P42 (GLMM p<0.05 & q<0.1). @ main effect of diet, &: interaction condition*diet, ^: significant difference with Tukey *post-hoc* test. p<0.05. Abbreviation: GLMM: General Linear Mixed Model.

Many changes were observed in the fecal microbiota composition of mice between P42 and P180 at all analyzed taxonomic levels (phylum, class, order, family, genus). At phylum level, the abundance of Bacteroidetes increased with age while Actinobacteria and Verrucomicrobia were found in lower abundances in P180 samples. At genus level, among many others, *Parasutterella* and *VadinBB60* increased and *RC9 gut group* and *Bilophila* decreased with age. All age-mediated changes and statistical vales are described in Figure 2 and Table 1.

**Table 1.**
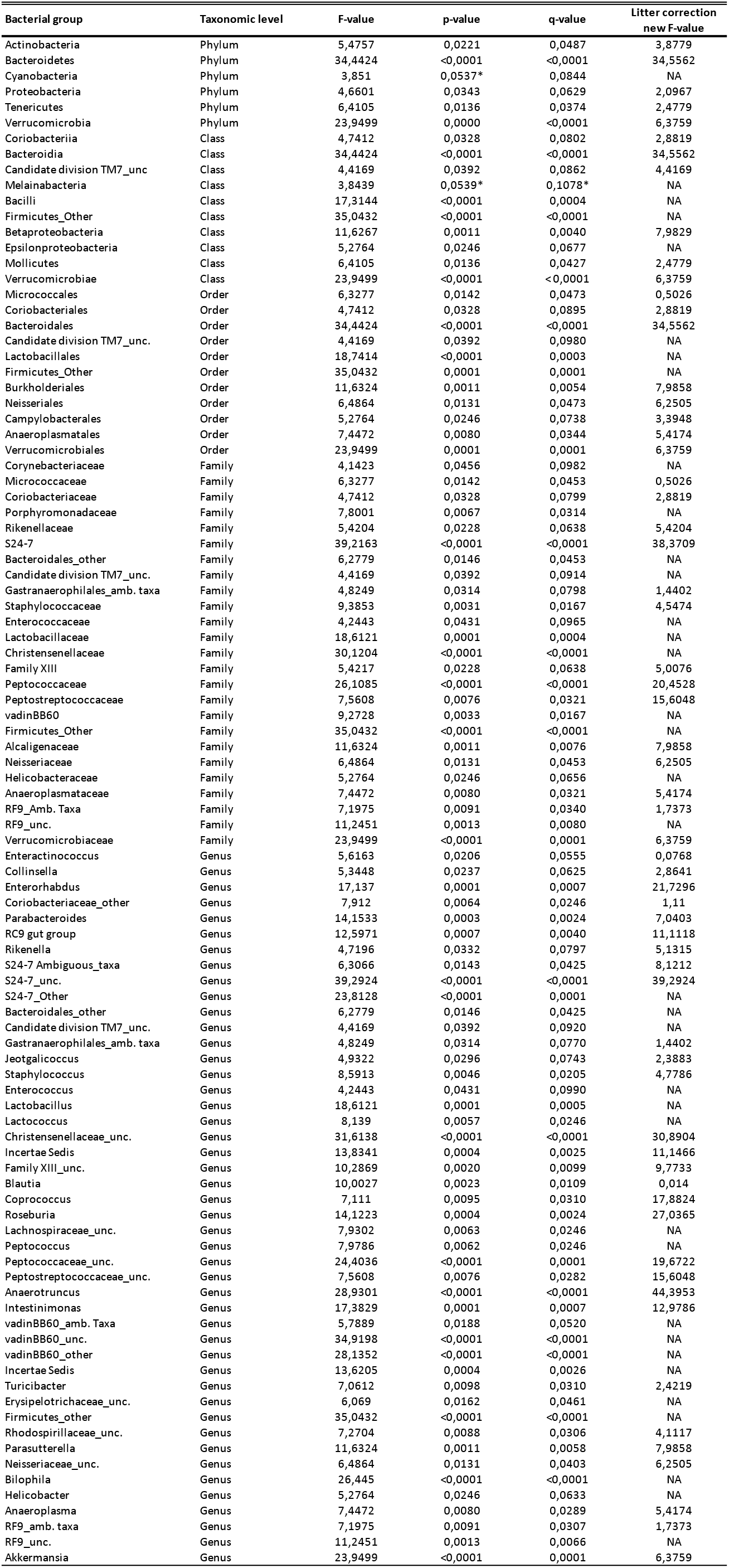
Significant age (P42 vs P180) effects on bacterial taxa at several taxonomic levels. GLMM p<0.05& q<0.1. *= trend.

Main effects for ELS and the early dietary ω-6/ω-3 ratio on the relative abundance were detected for bacterial groups at P42 and P180 (Fig. 3; Fig. 4; Table 2). At P42, ELS exposure decreased the abundance of *Coprococcus* and the low ω-6/ω-3 diet reduced the class, order and family Erysipelotrichia, Erysipelotrichales and Erysipelotrichaceae belonging to Firmicutes (Fig 3). At P180, the low ω-6/ω-3 diet long-lastingly reduced the genus *Coriobacteriaceae uncultured*. ELS reduced the relative abundance of the genera *RC9 gut group* and *Rikenella*, both part of the Rikenellaceae family in adulthood at P180 (Fig. 4).

**Figure 4.**
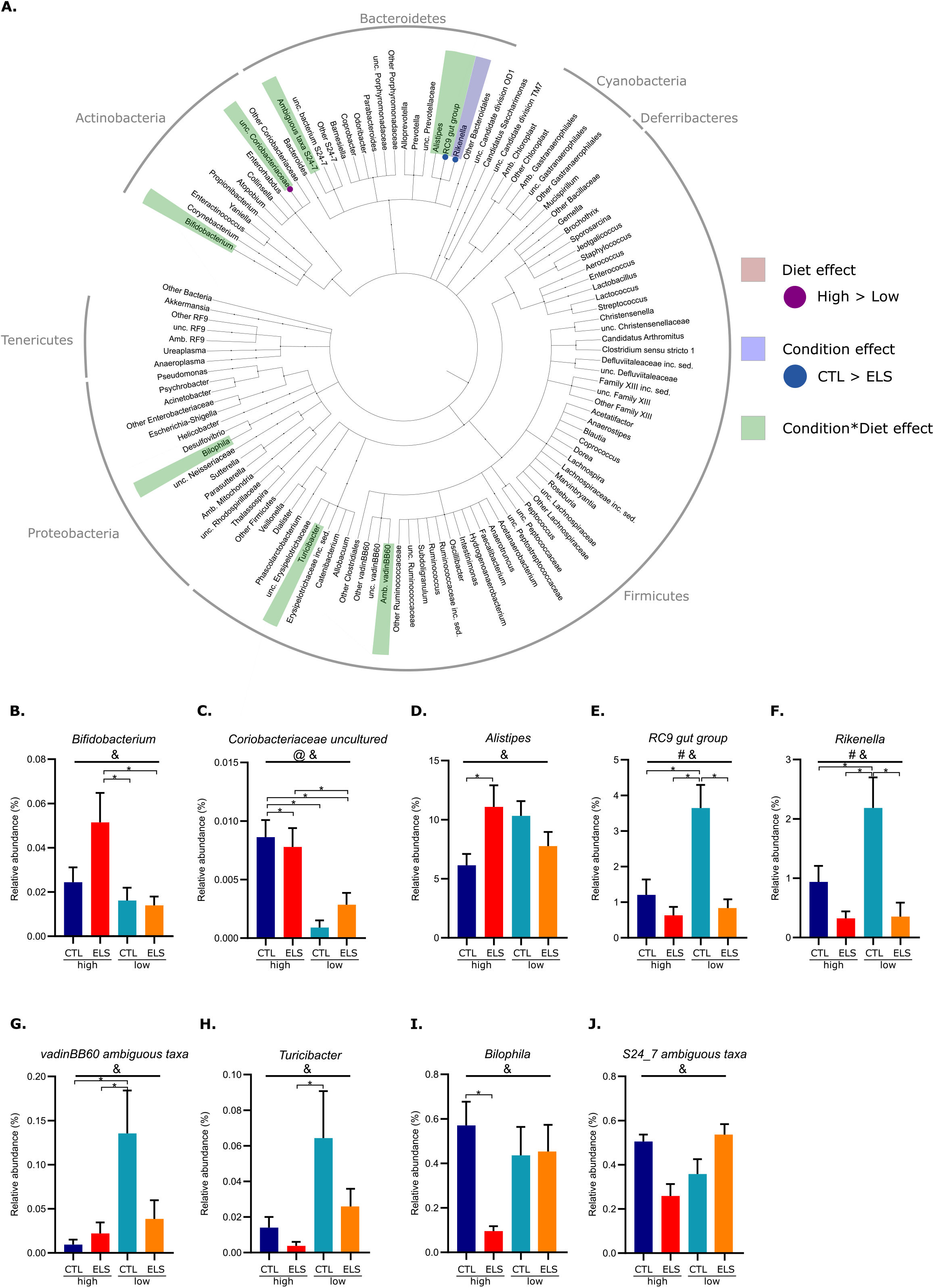
Early-life stress and early dietary ω-6/ω-3 ratio affect the microbiota composition at P180 in interaction with each other. **A:** Cladogram showing significant condition and diet-mediated changes in the relative abundance of bacterial taxa at several taxonomic levels at P180. **B - J:** Bar graphs of detected interaction effects (condition*diet) for bacterial taxa at P180 (GLMM p<0.05 & q<0.1). # main effect of Condition, @ main effect of diet, &: interaction condition*diet, ^: significant difference with Tukey *post-hoc* test. p<0.05. Abbreviation: GLMM: General Linear Mixed Model.

**Table 2.** Significant condition and diet effects on bacterial taxa at several taxonomic levels at P42 and P180. GLMM p<0.05& q<0.1.

| Bacterial group | Taxonomic level | Age | Effect | F-value | p-value | q-value | Litter correction |
| --- | --- | --- | --- | --- | --- | --- | --- |
|  |  |  |  |  |  |  | new F-value |
| Cyanobacteria | Phylum | P42 | Condition * Diet | 5.134 | 0.005 | 0.050 | NA |
| Melainabacteria | Class | P42 | Condition * Diet | 5.134 | 0.005 | 0.048 | NA |
| Gastranaerophilales | Order | P42 | Condition * Diet | 5.134 | 0.005 | 0.068 | NA |
| Erysipelotrichia | Class | P42 | Diet | 14.974 | 0.000 | 0.009 | 11.532 |
| Erysipelotrichia | Class | P42 | Condition * Diet | 6.277 | 0.002 | 0.033 | 5.057 |
| Erysipelotrichales | Order | P42 | Diet | 14.974 | 0.000 | 0.009 | 11.532 |
| Erysipelotrichales | Order | P42 | Condition * Diet | 6.277 | 0.002 | 0.047 | 5.057 |
| Erysipelotrichaceae | Family | P42 | Diet | 14.974 | 0.000 | 0.009 | 11.532 |
| Erysipelotrichaceae | Family | P42 | Condition * Diet | 6.277 | 0.002 | 0.044 | 5.057 |
| Allobaculum | Genus | P42 | Condition * Diet | 5.733 | 0.003 | 0.033 | NA |
| Clostridiales FamilyXIII | Family | P42 | Condition * Diet | 6.331 | 0.002 | 0.044 | NA |
| FamilyXIII IncertaeSedis | Genus | P42 | Condition * Diet | 6.931 | 0.001 | 0.026 | NA |
| FamilyXIII_unc. | Genus | P42 | Condition * Diet | 4.381 | 0.011 | 0.043 | NA |
| Clostridiales_Other | Family | P42 | Condition * Diet | 5.297 | 0.004 | 0.061 | NA |
| Clostridiales_Other | Genus | P42 | Condition * Diet | 5.297 | 0.004 | 0.033 | NA |
| Coprococcus | Genus | P42 | Condition | 2.985 | 0.045 | 0.148 | 4.302 |
| Odoribacter | Genus | P42 | Condition * Diet | 4.327 | 0.011 | 0.043 | 4.327 |
| Bifidobacterium | Genus | P180 | Condition * Diet | 4.027 | 0.014 | 0.036 | NA |
| Coriobacteriaceae_unc. | Genus | P180 | Diet | 24.924 | 0.000 | 0.001 | NA |
| Coriobacteriaceae_unc. | Genus | P180 | Condition * Diet | 8.575 | 0.000 | 0.002 | NA |
| Alistipes | Genus | P180 | Condition * Diet | 2.928 | 0.046 | 0.075 | NA |
| RC9 gut group | Genus | P180 | Condition | 10.670 | 0.002 | 0.074 | 4.950 |
| RC9 gut group | Genus | P180 | Condition * Diet | 10.437 | 0.000 | 0.001 | 6.522 |
| Rikenella | Genus | P180 | Condition | 12.487 | 0.001 | 0.070 | 8.791 |
| S24-7_Amb. taxa | Genus | P180 | Condition * Diet | 6.454 | 0.001 | 0.007 | 3.467 |
| VadinBB60_Amb. taxa | Genus | P180 | Condition * Diet | 4.677 | 0.007 | 0.026 | NA |
| Turicibacter | Genus | P180 | Condition * Diet | 3.601 | 0.022 | 0.043 | NA |
| Bilophila | Genus | P180 | Condition * Diet | 4.429 | 0.009 | 0.027 | NA |

At both ages, most significant changes in the relative abundance of the microbiota were dependent on both ELS and dietary ω-6/ω-3 ratio (Table 2). At P42 (Fig. 3), interaction effects were found between ELS exposure and diet for the phylum Cyanobacteria and its class and order Melainabacteria and Gastranaerophilales, for which the low diet significantly increased their abundance in specifically CTL animals, while in ELS animals no differences were present dependent on the early dietary ω-6/ω-3 ratio. This same pattern was found for several Clostridia members; Clostridiales Family XIII, an unassigned Clostridiales taxon*, Incertae sedis*, and an uncultured *Family XIII* taxon. Next, interaction effects between ELS and diet were detected for the class, order, family and genus Erysipelotrichia, Erysipelotrichales and Erysipelotrichaceae and *Allobaculum*, the low ω-6/ω-3 diet reduced its abundance in specifically CTL animals, while for ELS animals this reduction was not significant. Lastly, an interaction effect was found for the Bacteroidetes genus *Odoribacter*, for which ELS reduced its abundance in animals fed a high ω-6/ω-3 diet but not in animals fed the low ω-6/ω-3 diet.

At P180 (Fig. 4), an interaction between ELS and diet was found for the genus *Bifidobacterium*, relative abundance was significantly higher in ELS exposed animals fed the high ω-6/ω-3 diet as compared to CTL and ELS exposed animals fed the low ω-6/ω-3 diet. For the bacteria group *Coriobacteriaceae uncultured*, except from the reduction by the low ω-6/ω-3 diet for both CTL and ELS exposed animals as described above, an interaction between ELS exposure and diet was found. Next, an interaction effect was found for three members of the Rikenellaceae family. ELS exposure increased the abundance of *Alistipes* specifically in animals fed the high ω-6/ω-3 diet. For *RC9 gut group* and *Rikenella*, the ELS induced reduction (main effect ELS as described above), was only significant in animals fed the low ω-6/ω-3 diet. For the Firmicutes *VadinBB60 ambiguous taxa* and *Turicibacter* the low ω-6/ω-3 diet increased its abundance in CTL animals. Lastly, ELS exposure decreased the relative abundance of *Bilophila* only in animals fed the high ω-6/ω-3 diet. For *S24-7 ambiguous taxa* (Bacteroidetes) an interaction effect was found between ELS and diet at P180, however post hoc testing did not reveal significant differences between the experimental groups.

### 3.3. Correlations between bacterial taxa and peripheral and central outcome parameters within the same mice

We have recently reported that ELS exposure altered central and peripheral fatty acid profiles and impaired cognition in these animals^9^. Exposure to the low ω-6/ω-3 PUFA diet between P2 and P42 was able to protect against the ELS-induced cognitive deficits in adulthood but did not affect the metabolic alterations. In order to investigate if and how alterations in the microbiota might relate to these changes, we studied the correlation between several outcomes (behaviour, metabolic parameters and levels of central and peripheral fatty acid levels) and the relative abundance of bacterial groups at different taxonomic levels. All correlations are shown in Figure 5, and those with ρ>0.7 or ρ<-0.7 are reported in the text and supplementary Table S2 and S3.

**Figure 5.**
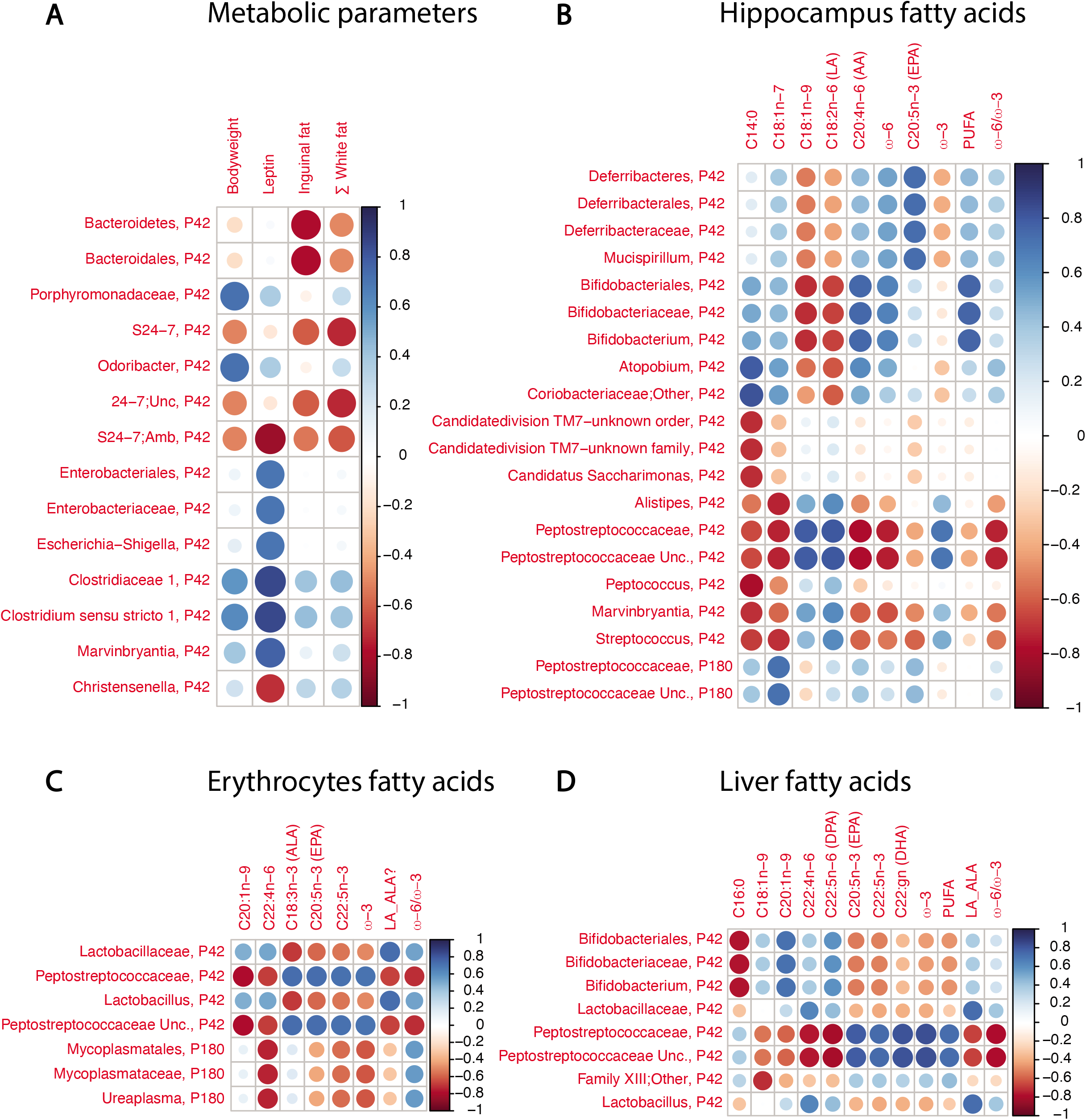
Bacterial taxa are correlated with several peripheral and central outcome parameters within the same mice. **A:** Correlations between bacterial taxa at P42 and metabolic outcomes parameters at P42. **B:** Correlations between bacterial taxa at P42 and fatty acid levels in the hippocampus at P42. **C:** Correlations between bacterial taxa at P42 are fatty acid levels in erythrocytes at P42. **D:** Correlations between bacterial taxa at P42 and fatty acid levels in the liver at P42. −1 < Spearman’s *rho* < 1

#### Bacterial taxa at P42 in relation to behaviour in adulthood

With regard to adult behaviour, we detected a negative correlation between the P42 levels of two related Bacteroidetes taxa *Porphyromonadaceae* and *Odoribacter* and performance on the object location task (OLT) (*rho=-0.7, rho = −0.73* respectively) (Supplementary Fig. 2). No correlations were detected for the other parameters related to behaviour.

#### Bacterial taxa at P42 in relation to metabolic outcome parameters at P42

The abundance of several bacterial species at P42 correlated with specific P42 metabolic outcomes (Fig. 5; Supplementary Table S2). Namely, the phylum Bacteroidetes and order Bacteroidales negatively correlated with the amount of inguinal fat (*rho* = −0.77 for both). Taxa of the Bacteroidetes phylum Porphyromonadaceae and *Odoribacter* positively correlated with bodyweight (*rho* = 0.73 for both). Several taxa within th*e* Proteobacteria phylum: Enterobacteriales, Enterobacteriaceae and *Escherichia-shigella* group (*rho* = 0.71), as well as taxa within the Firmicutes phylum: Clostridiaceae 1 and *Clostridium sensu stricto 1 (rho* = 0.85 for both) and *Marvinbryantia (rho* = 0.78) positively correlated with plasma leptin levels. The genus *Christensenella S24-7* negatively correlated with leptin levels (*rho* = −0.83 and −0.71 respectively). The Bacteroidetes S24-7 *and S24-7 Unc*. showed a negative correlation with the amount of white fat in mice (*rho* = −0.72 for both). There were no correlations between bacterial species at P180 and metabolic outcomes at P180.

#### Bacterial taxa at P42 in relation to fatty acid levels in the hippocampus, erythrocytes and liver at P42

We detected multiple strong correlations between bacterial taxa and fatty acid levels in the hippocampus, erythrocytes and liver (Fig. 5; Supplementary Table S3). From the Firmicutes phylum the Peptostreptococcaceae family negatively correlated with the ω-6/ω-3 ratio in the hippocampus, erythrocytes and liver (*rho* values of −0.76, −0.71, −0.77, respectively). In agreement with this, Peptostreptococcaceae positively correlated with ω-3 levels in all three tissues (*rho* > 0.7 for all). The Lactobacillaceae family, also from the Firmicutes phylum, positively correlated with the LA/ALA ratio in erythrocytes and liver (*rho* = 0.73 for both). Within the Actinobacteria phylum the *Bifidobacterium* lineage (from order, family until genus level) positively correlated with the amount of LCPUFAs in the hippocampus (*rho* = 0.82). We detected very few correlations between P180 bacterial species and fatty acid levels at P180 (Fig. 5; Supplementary Table S3)

## Discussion

We have previously shown that an early dietary intervention with reduced ω-6/ω-3 PUFA (LA/ALA) ratio protects against the ELS-induced cognitive deficits without affecting the metabolic alterations^9^. While the relation between stress, nutrition and the gut microbiota has been gaining increased attention over the recent years^13,32^, the specific mechanisms of such dietary interventions are not well understood. Here we demonstrate that chronic ELS during the first week of life (P2 - P9) increases the phylogenetic β-diversity of the gut microbiota both at P42 and persistently into adulthood (P180) in animals consuming a high ω-6/ω-3 diet. The early diet with low ω-6/ω-3 ratio was able to prevent this increase in β-diversity at P42, when animals were still consuming the experimental diet. In addition, ELS and the diet, mostly in interaction with each other, modulate the relative abundance of bacterial groups at several taxonomic levels on the short and long-term.

We will next discuss the microbiota diversity and composition across age, then elaborate on the short- and long-term impact of ELS and early dietary ω-6/ω-3 ratio on different microbiota parameters and lastly relate the microbial taxa abundance to earlier reported central and peripheral outcome measures from the same cohort of mice^9^.

### The microbiota across age

The phylogenetic diversity within samples (α-diversity) increased with age from weaning (P21) up to adulthood and, as expected, with only a relatively small difference between P42 and P180 samples in terms of the number of detected species. This is in line with previously described total amount of species across these ages^59,60^, while a decrease in α-diversity has been described in late adulthood or elderly which was associated with increased presence of diseases and medication^61^. The sample size at weaning age (P21) was relatively low, and even though the methodology that was used was reliable and sensitive enough to pick up age-related changes in α-diversity (sequencing depth of over 20.000 sequences for all three ages), we will further focus the discussion on our findings comparing adolescent (P42) and adult (P180) microbiota composition. The composition of the gut microbiota in terms of its relative species abundance is affected by age. Both at P42 and P180, Bacteroidetes and Firmicutes are the two most abundant phyla in all experimental groups, which is in line with other rodent and human microbiota profiles^62^. When comparing these two ages we observed multiple changes in the composition of the fecal microbiota, mainly consisting of a reduction in the phyla Actinobacteria and Verrucomicrobia (which includes the genus *Akkermansia*) and an increase of Bacteroidetes in adulthood. In particular, in P180 samples as compared to P42 samples, we observed lower abundance of members of the phylum Actinobacteria (Coriobacteriaceae and *Enterorhabdus*) of which *Bifidobacterium* is a genus and multiple members of the Firmicutes order Clostridiales. Also we observed higher abundances of members of the phylum Proteobacteria, such as the genus *Parasutterella*, in P180 samples as compared to P42 samples. In line with our comparative analyses between ages in mice, there is evidence for age dependent changes on the microbiome from human literature. While most studies up to date aimed at comparing gut microbiota of children between 0 and 2 years old with those of adults or elderly^63^ only very few have included adolescent groups. However, based on Agans et al. (2011), in line with our findings, adolescents can easily be separated from adults based on the relative species abundance and that in particular adolescent microbiota consist of a relative lower abundance of the genus *Sutterella* and relative higher abundance of *Bifidobacterium* and *Clostridium*^64,65^. Further work is needed, in both rodent and human cohorts, to be able to understand the age-related changes in microbiota composition in more detail and if and how each age group might be differently sensitive to stress exposure, diet or other environmental challenges.

### Short and long-term impact of early-life stress and early diet on microbiota composition

We will here first discuss the effects of ELS on microbiota α- and β-diversity and species abundance and thereafter the specific effects of the different dietary ω-6/ω-3 ratio on these parameters as well as the interaction of ω-6/ω-3 ratio with ELS exposure.

#### Short and long-term impact of early-life stress on microbiota composition

In this study, ELS exposure did not affect α-diversity at P42 or P180. Similar to our data, a multi-hit ELS model did not alter α-diversity in adult mice^66^, while our findings are in contrast with the ELS-reduction in α-diversity reported in rats, via the limited bedding and nesting (LBN) model at weaning^67^ or maternal separation (MS) in adulthood^68^. Thus, type of ELS model, outcome age and species seem to greatly impact the effects of early life adversity on the microbial α-diversity. In general, a less diverse microbiome is thought to be less resilient to external perturbations due to the loss of functional redundancy of the present species, therefore possibly less healthy^69^. However, whether health outcomes are positive or negative likely depends on the actual composition of the community.

The phylogenetic β-diversity distances between samples at OTU-level in ELS-exposed animals fed the high ω-6/ω-3-high diet, were strongly expanded both on short- and long-term, at P42 and P180. This suggests greater compositional differences between samples within the ELS group, meaning that the microbiota of ELS exposed mice are phylogenetically more apart from each other when compared to those of CTL mice. To our knowledge, this is the first time that such expansion of β-diversity is reported after ELS on the long-term and might suggests an aberrant microbial state, however the exact functional implications of such state remain to be understood^70,71^.

Few bacterial species were affected by ELS regardless of the early dietary ω-6/ω-3 ratio. At P42, ELS reduced the abundance of the genus *Coprococcus*, part of the Lachnospiraceae family. This is in line with the reduction in *Coprococcus* found at weaning in ELS exposed rats, via the LBN paradigm^67^. Lachnospiraceae and *Coprococcus* have been defined as major butyrate producing bacterial groups in both rodents and humans^72,73^, which suggests that ELS could affect butyrate levels via affecting these taxa. In adulthood (P180) the genera *RC9 gut group* and *Rikenella*, both part of the *Rikenellaceae* family and Bacteroidetes phylum were lastingly reduced by ELS. Similarly, MS in rats has been shown to reduce abundance of *Rikenella*, which also correlated with stress-induced corticosterone plasma levels in MS-exposed rats^42^. *Rikenella* is a well-known sugar fermenter and it has been suggested that stress can reduce the availability of sugars in the gut^74^ possibly leading to a decrease of bacteria involved in processing of sugars.

In summary, ELS during the first week of life does not affect α-diversity but leads to long-term effects on β-diversity. The expansion of the phylogenetic β-diversity between samples has been associated with an unhealthy or aberrant gut microbial state. Moreover, ELS affects the relative abundance of *Coprococcus* at P42 and *RC9* gut group and *Rikenella* at P180. The implications of these specific alterations within the relative abundance of bacterial groups are not well understood but nevertheless can impact the functionality of the gut microbiota.

#### Effect of the early dietary ω-6/ω-3 ratio and its interaction with early-life stress on the short and long term

We have previously reported, within this same cohort, a rescue effect of the low ω-6/ω-3 diet on the ELS-induced cognitive impairments as well as alterations in hippocampal brain plasticity, namely a reversal of the ELS-reduction in adult neurogenesis and the ELS-increase in the phagocytic marker of microglia, without affecting the ELS-mediated metabolic changes^9^. This allows us to not solely discuss effects of ELS and early diet on the microbiota, but also relate the observed changes to earlier described ELS-induced alterations, which were performed within the same mice cohort.

We found that the ELS-induced aberrant state of the phylogenetic β-diversity, the increased distances between animals fed a high ω-6/ω-3 diet early in life, was not present in animals that were consuming a low ω-6/ω-3 diet (as measured at P42). Because, as earlier described, the low ω-6/ω-3 diet protected against the ELS-induced central deficits in cognition and hippocampal plasticity in adulthood, it is tempting to speculate that this early modulation of β-diversity by dietary ω-6/ω-3 ratio could possibly modulate early developmental processes that contribute to the long-term ELS-induced brain-related outcomes^30,75^. Interestingly the effect of the diet on β-diversity is no longer present in adulthood. This is in line with the idea that diet mostly directly impacts microbiota. For example, it has been demonstrated in clinical trials that washout periods after dietary supplementation of DHA mostly revert DHA-mediated microbiota changes^76,77^ and that dietary patterns and fatty acid intake can be directly linked to the composition of the gut microbiota^78,79^. Similarly, in a preclinical setting, as mentioned before, it has been reported that life-long supplementation of ω-3 LCPUFAs can restore part of the disturbed gut microbiota composition of MS-exposed adult female rats, even though in this study no changes in β-diversity were reported^42^.

When looking at the relative abundance of microbial species, we see some immediate and long-lasting effects of the diet. Mice fed the low ω-6/ω-3 ratio diet from P2-P42, exhibited a reduction in Erysipelotrichia lineage down to the Erysipelotrichaceae family (Firmicutes) when compared to mice fed the high ω-6/ω-3 ratio diet. These taxa have been reported to be increased in obese individuals notoriously consuming diets with excess of ω-6 fatty acids^80^, pointing towards the idea that dietary ω-6/ω-3 ratio is an important modulator of these specific bacteria and their balance. Similarly, ω-3 (LC)PUFA supplementation has been shown to lead to a decrease of the Firmicutes phylum^33,81,82^ and restoration of the Firmicutes/Bacteroidetes ratio, often reported to be higher under pathological conditions such as obesity and inflammatory bowel syndrome (IBS)^83,84^.

Next to the independent effects of ELS and dietary ω-6/ω-3 ratio, their interaction is particularly interesting to gain further insight in how the diet might exert its protective effect on the ELS-induced deficits. For example, directly after the end of the dietary intervention at P42, specifically control mice fed the low ω-6/ω-3 diet exhibited an increased abundance of several Clostridia members when compared to those fed the diet with the high ω-6/ω-3 ratio. These taxa belong to the phylum Firmicutes and order Clostridiales that are known for their involvement in the production of butyrate^85,86^. Such modulation is in line with the fact that ω-3 fatty acid supplementation can indeed lead to increased bacterial derived butyrate^72,77^. Short-chain fatty acids (SCFA) such as butyrate, propionate and acetate are bacterial-derived metabolites of fibers and carbohydrate that have been suggested to be key for mental health^87^. For example by increasing central brain derived neurotrophic factor (BDNF) production^88,89^ and modulation of microglial maturation and functionality^90^. As mentioned above, the low ω-6/ω-3 diet was able to prevent ELS-induced alterations in hippocampal plasticity including microglial morphology and phagocytic capacity^9^, raising the question if this could possibly be related to an increase in bacterial-derived butyrate by the diet in ELS exposed animals specifically. Next, ELS exposure reduced the abundance of the Bacteroidetes genus *Odoribacter* in animals fed the high ω-6/ω-3 diet but not in animals fed the low ω-6/ω-3 diet. *Odoribacter* is a known producer of acetate, propionate and butyrate^91^, decreased *Odoribacter* may affect host inflammation via reduced SCFA availability. It will be interesting to see in follow-up studies whether indeed our conditions lead to an altered levels of SCFAs and in particular butyrate.

Summarizing, we found that low dietary ω-6/ω-3 ratio prevents the ELS-induced expansion of the phylogenetic β-diversity. In addition, the dietary ω-6/ω-3 ratio significantly impacted the presence of microbes in the gut while animals were still consuming the experimental diet, mostly in interaction with ELS exposure. Many of these changes are in line with literature showing beneficial effects of dietary ω-3 supplementation on brain and metabolism^40,92^ and suggest that the diet induced protective effects might be partly modulated by the observed changes in microbiota. We hypothesize that lowering ω-6/ω-3 early in life, via lowering dietary LA/ALA ratio, contributes to a stable and diverse microbiota thereby affecting sensitive developmental processes that could impact the later-life health status.

Important to note is that within the current study, all animals were on a life-long synthetic diet, enabling us to control for the source and proportion of its ingredients. Such synthetic diets, also referred to as “refined”, contain mostly insoluble fibers such as cellulose (Supplementary Table S4), which distinguishes it from the regular chow diets containing both soluble and insoluble fibers. These dietary conditions likely impact microbiota composition since distinct bacterial species are involved in the fermentation of soluble versus insoluble fibers^93,94^. While this does not affect the differences observed between groups in the current study as all experimental groups were exposed to synthetic diet, it is important to bare this in mind when comparing the current findings to existing literature^95^.

#### Abundance of microbiota species in relation to central and peripheral outcomes

We studied how the bacterial changes correlate with the previously published ELS- and diet-mediated differences in cognitive abilities, metabolic alterations and central and peripheral fatty acid profiles analyzed in this same cohort^9^. As mentioned in section *“Short and long-term impact of early-life stress on microbiota composition*” we found a negative correlation between *adult* performance on a spatial memory task and the levels of Bacteroidetes family *Porphyromonadaceae* and its genus *Odoribacter at P42* and not at P180. Similarly, an increased abundance in both taxa have been described in aged mice^96^ and specifically *Porphyromonadaceae* has been shown to be negatively correlated with cognitive dysfunction in humans^97,98^. Suggesting that a dysregulation of these taxa might be key in modulating cognitive functions. We have previously reported that ELS exposure leads to a life-long reduction in white fat mass and circulating leptin^99^, while these ELS-induced effects were not modulated by the diet^9^. When studying the correlation of the bacterial profile with the metabolic outcomes (body weight, fat mass and leptin) we found a positive correlation of Porphyromonadaceae and *Odoribacter* with bodyweight at P42, which is in line with a previous report showing that their abundance is increased in HFD-exposed mice^100^. In addition, the phylum Bacteroidetes and multiple of its taxa (e.g. *S24-7*), were negatively correlated with the amount of white fat mass. Notably, an unidentified taxon from the S24-7 family has been reported to be affected by early life supplementation of synbiotics that protected against diet-induced obesity in adult mice^101^. Indeed high levels of Bacteroidetes and some of its taxa are associated with a healthy non-Western diet while lower levels are associated with a Western-style diet^32,102^. Finally, there was a positive correlation between several taxa within the Proteobacteria phylum and plasma leptin levels. Importantly, changes in the Proteobacteria have been associated with HFD in mice and humans, where leptin levels are dysregulated as well^79,80,103^. Also, Bacteroidetes *S24-7-ambiguous taxa* and the Firmicutes genus *Christensenella* were negatively correlated with plasma leptin, both associated with reduction in body weight or adiposity in mice^101,104^, suggesting that these bacteria might be particularly sensitive to conditions with altered leptin and fat mass. Lastly, we found multiple strong correlations between bacterial species and specific fatty acid levels in the hippocampus, liver and erythrocytes (Fig. 5; Supplementary Table S3). To name a few examples, there was a negative correlation between the P42 hippocampal, liver and erythrocyte ω-6/ω-3 ratio and the relative abundance of Peptostreptococcaceae (Firmicutes) at P42. In agreement, Peptostreptococcaceae positively correlated with ω-3 levels in all three tissues. Interestingly a life-long ω-3 PUFA supplementation starting prenatally lead to decreased levels of Peptostreptococcaceae when compared to chow-fed or ω-3 deficient mice^105^. Such discrepancy is mostly likely due to the length and type of the dietary intervention. Next, the *Bifidobacteria*, which has been established to be increased by diets high in ω-3 fatty acids^77,105^, correlated with the amount of hippocampal PUFAs. Several positive functions have been attributed to *Bifidobacteria* such as degradation of non-digestible carbohydrates, production of vitamin B, antioxidants, stimulation of the immune system, and increasing butyrate levels via cross-feeding^106,107^. While the above-mentioned relations are of course of descriptive nature, they give us a lead for future investigations to better understand which processes might be most impacted by microbiota changes and via which routes the microbiota changes could be involved in the observed ELS and diet induced effects.

In conclusion, we show that exposure to ELS via the LBN paradigm during the first postnatal week and the ω-6/ω-3 ratio of the early diet from P2-P42 affect the gut microbiota of male mice. These data give novel insights in the complex interaction between ELS, early dietary ω-3 availability and the gut microbiota across ages and provide a basis for i) future studies addressing the causal relationship between the alterations in microbiota, the ELS-induced deficits and diet ii) as well as for non-invasive (nutritional) interventions targeting the microbiota to protect against and/or reverse the ELS-induced deficits.

## Data availability statement

The data that support the findings of this study are openly available in “figshare” at https://doi.org/10.6084/m9.figshare.16748824.v1

## Author contributions

KR analyzed the data, prepared the figures and wrote the manuscript. ST analyzed the data and prepared the figures. KY, LS and AK conceptualized the study and KY performed the mouse-related experimental work. MM contributed to correlation analysis and discussion interpretation. AK supervised this study and reviewed and edited the manuscript. All authors contributed to editing of the manuscript.

## Acknowledgement

We would like to thank H. de Weerd for her contribution to the statistical analysis and P. van Leeuwen for his assistance with interpretation of the data.

---

**Supplementary figure 1. Age impacts phylogenetic β-diversity**

**A, B:** Principal Component Analysis (PCA) and distance based Redundancy Analysis (db-RDA) of β-diversity aggregated at genus level for both ages. **C, D**: Average weighted UniFrac distances (between groups) of β-diversity on OTU level comparing phylogenetic configurations of fecal microbial communities of the four experimental groups for both ages.

**Supplementary figure 2. Bacteroidetes taxa *Porphyromonadaceae* and *Odoribacter* negatively correlate with performance on the object location task (OLT)**

